## Supplemental material for "Meiotic DNA break resection and recombination rely on chromatin remodeler Fun30"

**This pdf contains:**

Supplementary Figures 1–7
Supplementary Tables 1 and 2.

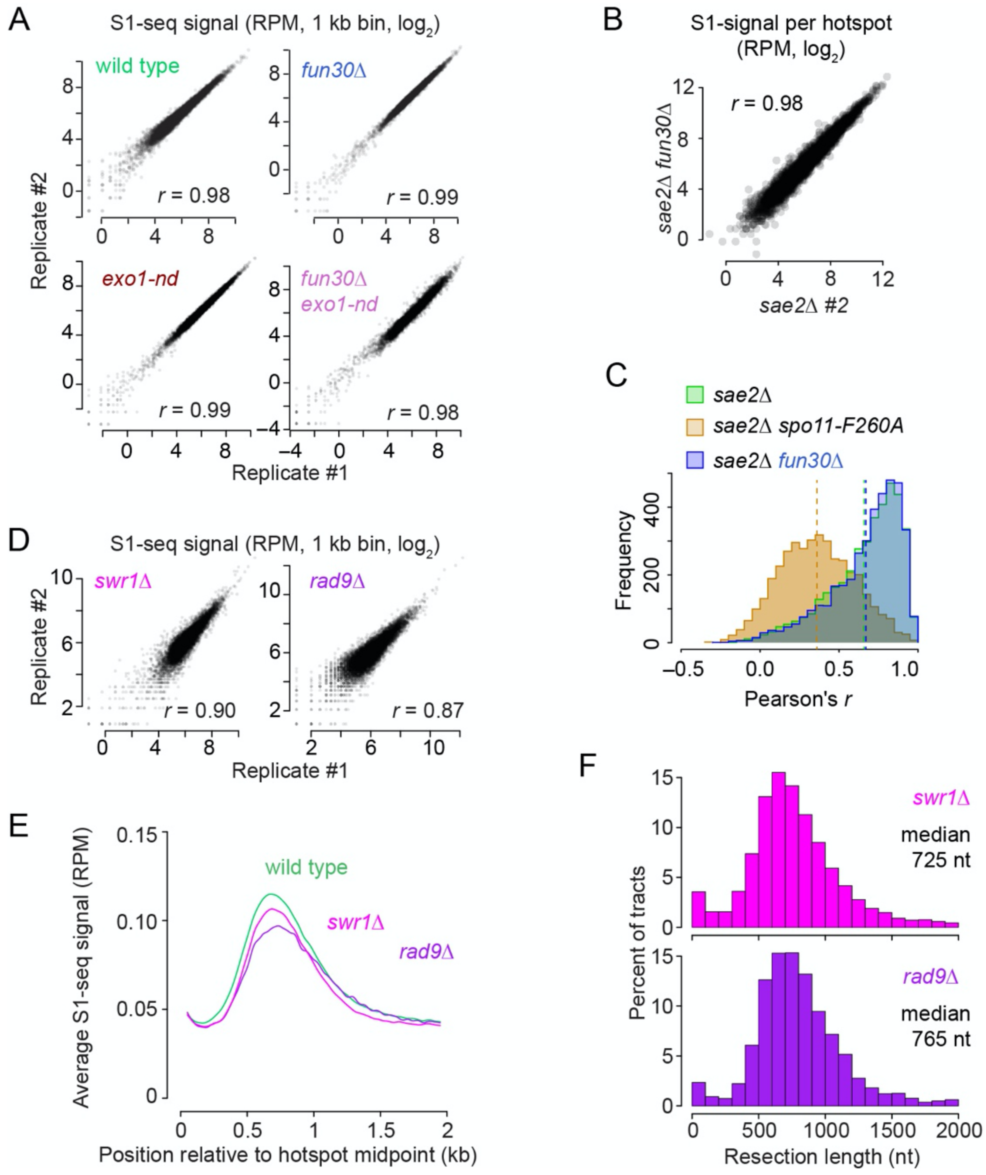


**Figure S1. S1-seq reproducibility and S1-seq in *swr1*Δ and *rad9*Δ**

(A) Reproducibility of S1-seq between two biological replicates of each genotype. Each point is the $\log_2$-transformed read count in a 1-kb segment of the genome.

(B) Preservation of hotspot heats in *fun30*Δ. Each dot represents the sum of the S1-seq signal at a given hotspot in a *sae2*Δ background (n = 3908). S1-seq library preparation was previously shown to quantitatively capture the unresected DSBs that accumulate in *sae2*Δ mutants (Mimitou et al. 2017; Mimitou and Keeney 2018).

(C) Preservation of fine-scale DSB distributions within hotspots in *fun30*Δ. For each of 3908 DSB hotspots, we computed the correlation coefficient (Pearson's *r*) comparing the S1-seq spatial distribution between the indicated pairs of datasets, then plotted the distributions of the 3908 correlation coefficients. The vertical dashed lines indicate the mean *r* value for each comparison. We compared two biological replicates of *sae2*Δ to show the intrinsic variability in this measurement (mean *r* = 0.66) and we compared *sae2*Δ to *sae2*Δ *spo11-F260A* (Claeys Bouuaert et al. 2021) as an example of what happens in a mutant with altered DSB distributions (mean *r* = 0.36). Comparison of *sae2*Δ with *fun30*Δ *sae2*Δ gave a distribution of correlation coefficients indistinguishable from the comparison of *sae2*Δ replicates (mean *r* = 0.67), indicating that the *fun30*Δ mutation has little or no measurable effect on local DSB distributions within hotspots.

(D) Reproducibility of *swr1*Δ and *rad9*Δ S1-seq biological replicates.

(E) Average S1-seq distribution around hotspots in *swr1*Δ and *rad9*Δ*.*

(F) Distribution of resection tract lengths in *swr1*Δ and *rad9*Δ as in **Figure 2C**.

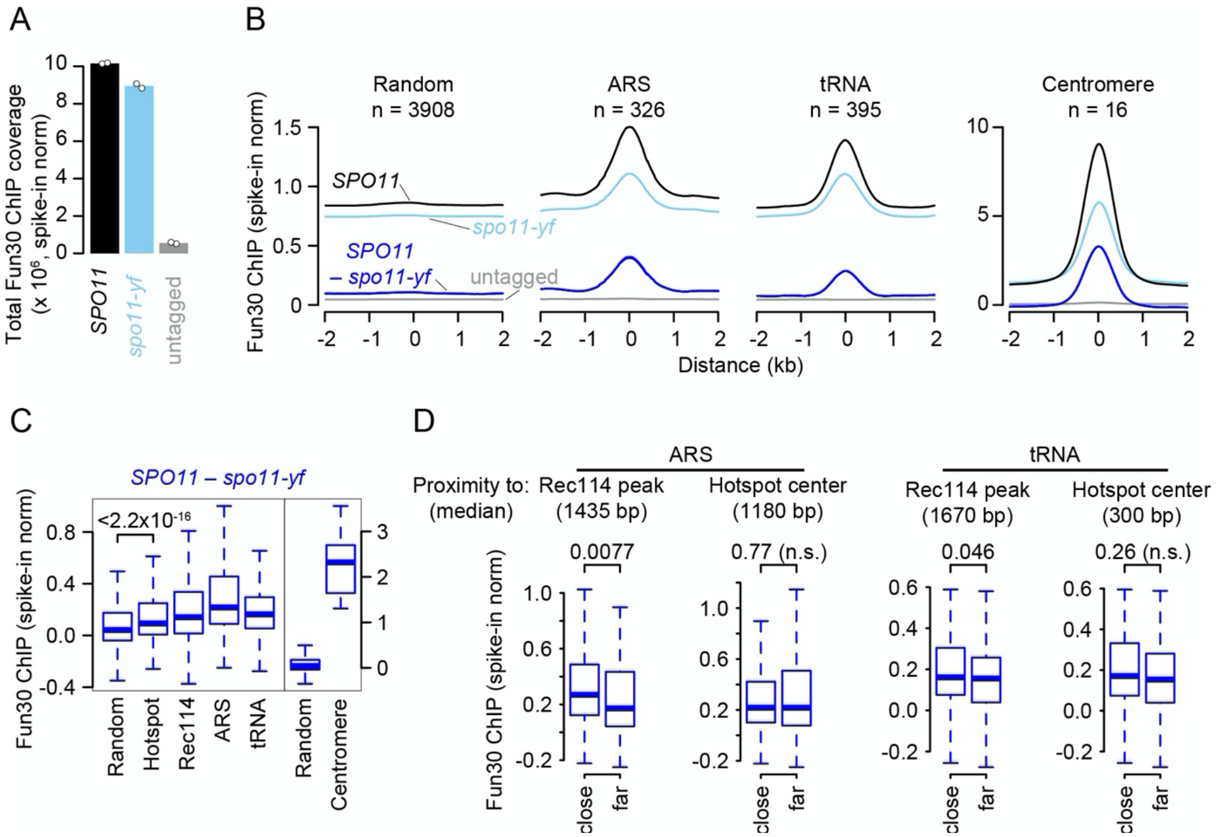


**Figure S2. DSB-dependent Fun30 enrichment**

(A) Total Fun30 ChIP-seq coverage normalized to the spike-in control. Bars are the means from two biological replicates; open circles show the individual values for each replicate.

(B) Average Fun30 ChIP-seq signals around ARS, tRNA, and centromere. The random sites here and in panel C are the same as in **Figure 3B**.

(C) DSB-dependent Fun30 enrichment. Box plots summarize the distributions across all of the indicated elements from panel B and **Figure 3B** for Fun30 ChIP-seq signal summed in 1-kb windows. Note the different y-axis scales for left and right parts of the plot. In all box plots, thick horizontal bars denote medians, box edges mark the upper and lower quartiles, and whiskers indicate values within 1.5-fold of the interquartile range. Outliers are not shown. Here and in panel D, numbers above brackets indicate *P* values of two-sided Wilcoxon tests.

(D) ARS and tRNAs that are closer to Rec114 binding sites tend to exhibit higher DSB-dependent Fun30 ChIP-seq signals. The ARS and tRNA regions from panels B and C were subdivided into two groups based on the distance to the nearest Rec114 peak or hotspot center: “close” indicates elements less than the median distance away and “far” indicates the rest. Proximity to Rec114 peaks was associated with a significantly higher DSB-dependent Fun30 ChIP-seq signal, whereas proximity to hotspots showed no such pattern. These results suggest that at least some of the DSB-dependent recruitment of Fun30 to ARS or tRNA genes is a consequence of fortuitous proximity or overlap of (some of) these elements with Rec114 ChIP peaks.

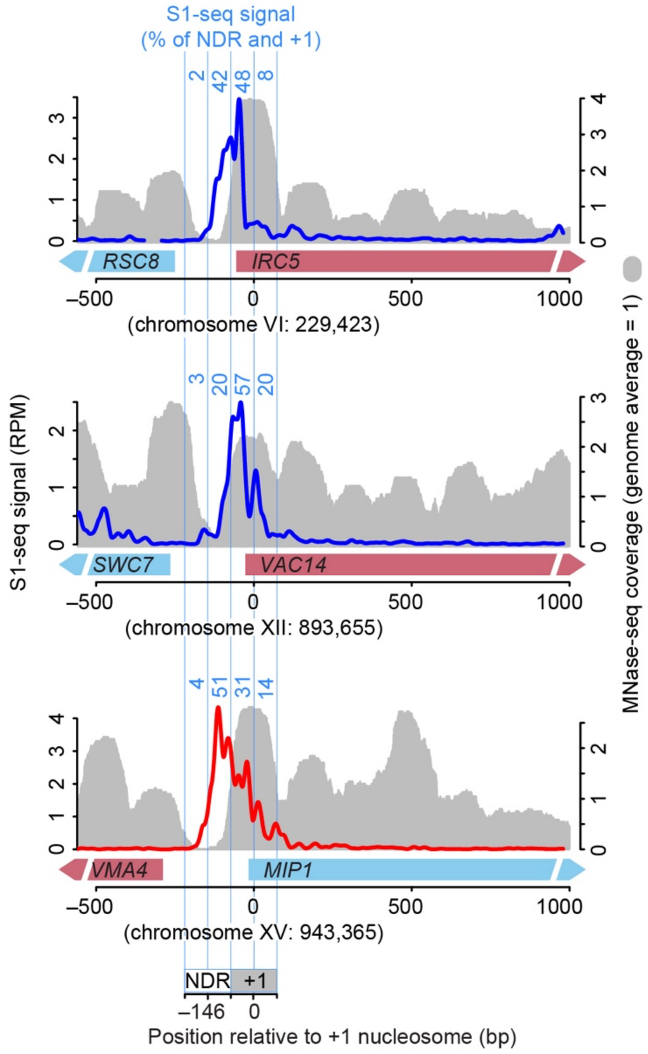


**Figure S3. MRX/Sae2 nicks within the +1 nucleosome**
Examples of resection endpoint distributions in *exo1-nd fun30*Δ at three representative loci that contributed to the average shown in **Figure 4B**. S1-seq signals (41-bp smoothed) from the top (blue) or bottom (red) strand are shown, dependent on the orientation of the gene where the +1 nucleosome is located. Numbers in light blue indicate the percentages of S1-seq signal in the four windows spanning the +1 nucleosome and NDR (see legend to **Figure 4B**).

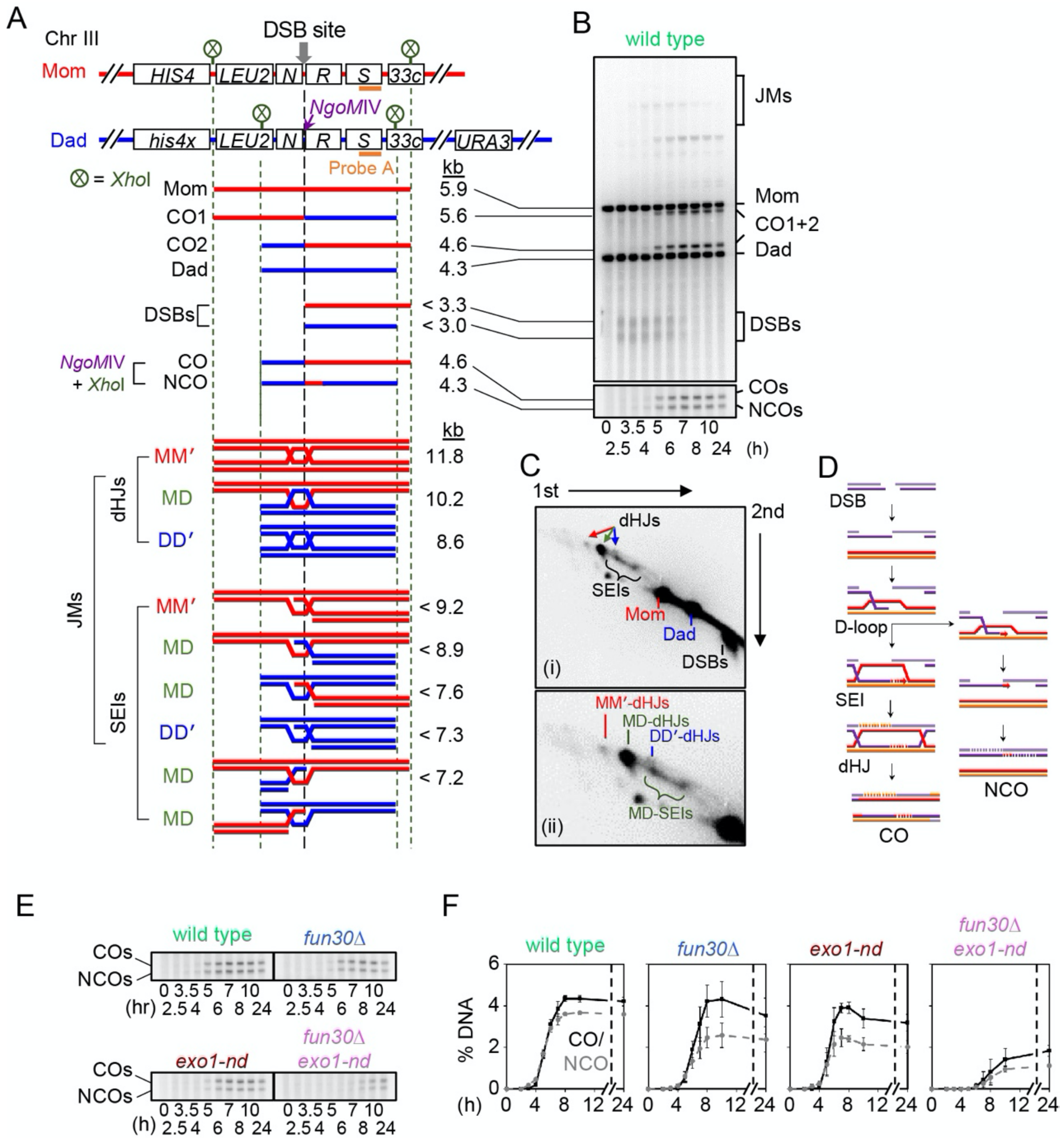


**Figure S4. Physical assay detecting recombination intermediates at the *HIS4LEU2* hotspot**

(A) Physical map of the *HIS4LEU2* locus showing diagnostic *Xho*I restriction enzyme sites and the position of Southern blot probe A. “Mom” and “Dad” indicate the two parental versions of the locus; COs, crossovers; NCOs, non-crossovers; MM' IS-dHJ, intersister double-Holliday junction; MD IH-dHJ, interhomolog double-Holliday junction; DD' IS-dHJ, intersister double-Holliday junction; SEIs, single-end invasions. Positions of *Xho*I sites are indicated as circled Xs.

(B) Example one-dimensional gel analysis showing parental signals, DSBs, COs, NCOs, and joint molecules (JMs). The Southern blot image is reproduced from **Figure 6A**.

(C) Example two-dimensional gel displaying parental signals and recombination intermediates. Green arrow or text indicate interhomolog species; red and blue arrows and text indicate intersister species.

(D) Key steps in crossover and noncrossover formation during meiosis.

(E) Representative gel images of crossovers and noncrossovers.

(F) Quantification of crossovers and noncrossovers (mean ± SD for three independent meiotic cultures).

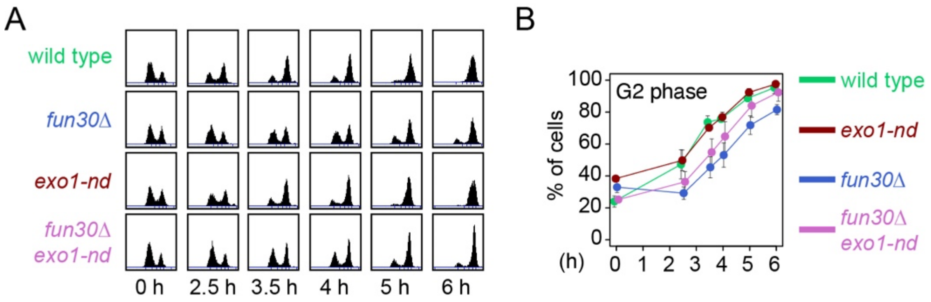


**Figure S5. Analysis of early meiotic progression by flow cytometric DNA analysis.**
(A) Representative histograms of flow cytometric measurements of the cellular DNA content.
(B) Progression of pre-meiotic DNA replication during early meiosis. The percentage of cells in G2 phase was calculated from flow cytometry (mean ± SD for three independent meiotic cultures).

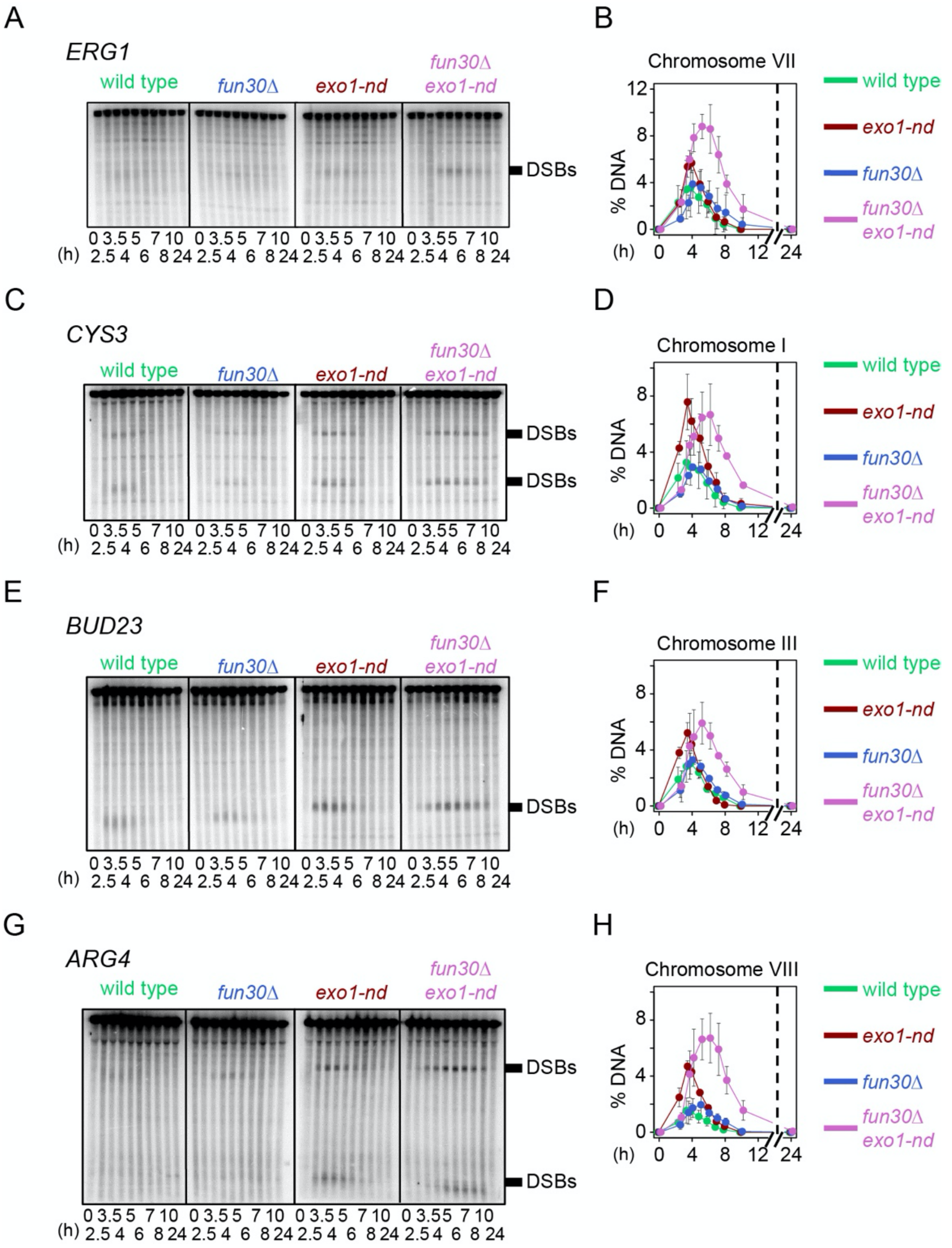


**Figure S6. DSBs formation at various natural hotspots.**
(A–H) Representative one-dimensional gel analyses of DSBs and corresponding quantification at the *ERG1* (**A and B**), *CYS3* (**C and D**), *BUD23* (**E and F**), and *ARG4* (**G and H**) hotspots. Error bars indicate mean ± SD for three independent cultures.

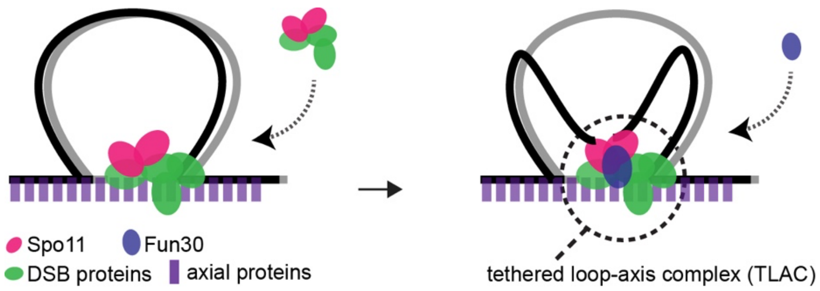


**Figure S7. Schematic representation of DSB-dependent Fun30 recruitment in the tethered loop-axis complex**

A model proposed based on findings in this study. In response to Spo11 cleaving DNA within the TALC, Fun30 is recruited to the DSB ends and remodels the nucleosomes.

**Table S1. List of yeast strains used in this study.**

| S1-seq strains | strains | MAT | Genotypes |
|---|---|---|---|
| Wild type | SKY6066 | *a/α* | *nuc1Δ::LEU2/"* |
| *exo1-nd* | SKY 6075 | *a/α* | *nuc1Δ::LEU2/", exo1-nd/"* |
| *fun30Δ* | SKY *6457* | *a/α* | *nuc1Δ::LEU2/", fun30Δ::KanMX/"* |
| *fun30Δ exo1-nd* | SKY 6593 | *a/α* | *nuc1Δ::LEU2/", fun30Δ::KanMX/", exo1-nd/"* |
| *sae2Δ* | SKY 6436 | *a/α* | *nuc1Δ::LEU2/", sae2Δ::NatMX/"* |
| *fun30Δ sae2Δ* | SKY 6587 | *a/α* | *nuc1Δ::LEU2/", fun30Δ::KanMX/", sae2Δ::NatMX/"* |
| S1 Southern and S1-seq strains | strains | MAT | genotypes |
| WT | SKY 6066 | *a/α* | *nuc1Δ::LEU2/"* |
| *sae2Δ* | SKY 6436 | *a/Tα* | *nuc1Δ::LEU2/", sae2Δ::NatMX/"* |
| *exo1-nd* | SKY 6075 | *a/α* | *nuc1Δ::LEU2/", exo1-nd/"* |
| *arp8Δ* | SKY 6094 | *a/α* | *nuc1Δ::LEU2/", arp8Δ::KanMX4/"* |
| *htz1Δ* | SKY 6081 | *a/α* | *nuc1Δ::LEU2/", htz1Δ::KanMX/"* |
| *swr1Δ* | SKY 6430 | *a/α* | *nuc1Δ::LEU2/", swr1Δ::KanMX/"* |
| *fun30Δ* | SKY 6457 | *a/α* | *nuc1Δ::LEU2/", fun30Δ::KanMX/"* |
| *fun30Δ sae2Δ* | SKY 6587 | *a/α* | *nuc1Δ::LEU2/", fun30Δ::KanMX/", sae2Δ::NatMX/"* |
| *fun30Δ exo1-nd* | SKY 6593 | *a/α* | *nuc1Δ::LEU2/", fun30Δ::KanMX/", exo1-nd/"* |
| *fun30k603R* | SKY 6736 | *a/α* | *nuc1Δ::LEU2/", fun30K603R::URA3/", fun30Δ::KanMX/"* |
| *arp8Δ htz1Δ* | SKY 6424 | *a/α* | *nuc1Δ::LEU2/", arp8Δ::KanMX/", htz1Δ::KanMX4/"* |
| *arp8Δ swr1Δ* | SKY 6504 | *a/α* | *nuc1Δ::LEU2/", arp8Δ::KanMX/", swr1Δ::KanMX4/"* |
| *arp8Δ fun30Δ* | SKY 6498 | *a/α* | *nuc1Δ::LEU2/", arp8Δ::KanMX/", fun30Δ::KanMX4/"* |
| *htz1Δ swr1Δ* | SKY 6492 | *a/α* | *nuc1Δ::LEU2/", htz1Δ::KanMX/", swr1Δ::KanMX4/"* |
| *htz1Δ fun30Δ* | SKY 6486 | *a/α* | *nuc1Δ::LEU2/", htz1Δ::KanMX/", fun30Δ::KanMX4/"* |
| *swr1Δ fun30Δ* | SKY 6510 | *a/α* | *nuc1Δ::LEU2/", swr1Δ::KanMX/", fun30Δ::KanMX4/"* |
| Flourescent spore assay (FSA) | strains | MAT | genotypes |
| *THR1::m-Cerulean* | SKY 3576 | *a* | *trp1::hisG, THR1::m-Cerulean-TRP1* |
| *CEN8::tdTomato-, ARG4::GFP** | SKY 3579 | *α* | *trp1::hisG, CEN8::tdTomato-LEU2, ARG4::GFP*-URA3* |
| *fun30Δ exo1-nd_THR1::m-Cerulean* | SKY 6779 | *a* | *trp1::hisG, fun30Δ::KanMX, THR1::m-Cerulean-TRP1, exo1-nd* |
| *fun30Δ exo1-nd_CEN8::tdTomato-, ARG4::GFP** | SKY 6774 | *α* | *trp1::hisG, fun30Δ::KanMX, CEN8::tdTomato-LEU2, ARG4::GFP*-URA3, exo1-nd* |
| ChIP-seq | strains | MAT | genotypes |
| untagged | SKY 6066 | *a/α* | *nuc1Δ::LEU2/"* |
| *Fun30-myc sae2Δ* | SKY 6822 | *a/α* | *sae2Δ::NatMX/", nuc1Δ::LEU2/", Fun30-myc-KanMX/"* |
| *fun30K603R-myc sae2Δ* | SKY 7085 | *a/α* | *nuc1Δ::LEU2/", fun30K603R-13myc-KanMX6/", sae2Δ::NAT/"* |
| *Fun30-myc spo11-yf sae2Δ* | SKY 7231 | *a/α* | *ho::LYS2/", lys2/", ura3/", leu2::hisG/", spo11yf::KanMX4/", sae2Δ::NatMX/", nuc1Δ::LEU2/", Fun30-myc-KanMX/"* |
| *Myc-tagged Rec114 (S.mikatae IFO1815): spike in* | SKY 6052 | *a/α* | *REC114-myc8::ura3::HphMX6/REC114-myc8::ura3::HphMX6* |
| Physical analysis | strains† | MAT | *genotypes‡* |
| | KKY2945 | *a/α* | *HIS4::LEU2-(BamHI)/his4x::LEU2-(NgoMIV)--URA3, nuc1Δ::LEU2, ERG1::SalI/ERG1::SpeI* |
| | KKY6069 | *a/α* | *HIS4::LEU2-(BamHI)/his4x::LEU2-(NgoMIV)--URA3, nuc1Δ::LEU2, ERG1::SalI/ERG1::SpeI, fun30Δ::KanMX4* |
| | KKY6101 | *a/α* | *HIS4::LEU2-(BamHI)/his4x::LEU2-(NgoMIV)--URA3, nuc1Δ::LEU2, ERG1::SalI/ERG1::SpeI, exo1(D173A)* |
| | KKY4555 | *a/α* | *HIS4::LEU2-(BamHI)/his4x::LEU2-(NgoMIV)--URA3, nuc1Δ::LEU2, ERG1::SalI/ERG1::SpeI, fun30::KanMX4, exo1(D173A)* |

† All strains are isogenic derivatives of SK1 background.

‡ All strains are also homozygous for the mutation *ho::hisG*, *leu2::hisG*, *ura3 (ΔPstI-SmaI)*.

**Table S2. Genetic distance and MI nondisjunction estimated by fluorescent spore assay**

| | | wild type | *fun30Δ exo1-nd* |
|---|---|---|---|
| *CEN8-ARG4* | PD:TT:NPD | 720:256:0 | 1265:332:2 |
| | cM ± SE | 13.11 ± 0.7 | 10.76 ± 0.57 |
| | *p* (vs wild type) | N/A | 0.00253 |
| *ARG4-THR1* | PD:TT:NPD | 884:92:0 | 1430:169:0 |
| | cM ± SE | 4.71 ± 0.47 | 5.28 ± 0.38 |
| | *p* (vs wild type) | N/A | 0.645 |
| | MI-NDJ | 3/979 = 0.3% | 37/1636 = 2.26% |

The number of tetrads exhibiting fluorescent marker configurations in parental ditype (PD), tetratype (TT) and nonparental ditype (NPD) are shown. Genetic distances were calculated using Perkins equation [cM = 100*(6*NPD+TT)/(2*(PD+NPD+TT))]. Standard errors (SE) were calculated using Stahl Lab Online Tools (https://elizabethhousworth.com/StahlLabOnlineTools/). NPD in the *ARG4–THR1* interval was omitted as they are indistinguishable with MI-NDJ (Thacker et al. 2011). Wild type data are from our previous publication (Thacker et al. 2011). *p* values for genetic distance were calculated by G test.

**Table S3. Summary of genomic datasets**

**S1-seq data (GSE221377)**

| | |
|---|---|
| s_sky6066_4h_index1_HITMAP | wild type |
| s_sky6066_4h_index2_HITMAP | wild type |
| s_sky6075_4h_index5_HITMAP | *exo1-nd* |
| s_sky6075_4h_index6_HITMAP | *exo1-nd* |
| s_sky6057_4h_index1_HITMAP | *fun30Δ* |
| s_sky6057_4h_index2_HITMAP | *fun30Δ* |
| s_sky6593_4h_index3_HITMAP | *fun30Δ exo1-nd* |
| s_sky6593_4h_index4_HITMAP | *fun30Δ exo1-nd* |
| s_SKY6436_4h_1_HITMAP | *sae2Δ* |
| s_SKY6587_4h_1_HITMAP | *fun30Δ sae2Δ* |
| s_SKY6587_4h_2_HITMAP | *fun30Δ sae2Δ* |

**ChIP-seq data (GSE221033)**

| | |
|---|---|
| Project#11182 | |
| s_sky6066-1input | untagged |
| s_sky6066-2input | untagged |
| s_sky6822-1input | *Fun30myc sae2Δ* |
| s_sky6822-2input | *Fun30myc sae2Δ* |
| s_sky7231-1input | *Fun30myc spo11yf sae2Δ* |
| s_sky7231-2input | *Fun30myc spo11yf sae2Δ* |
| s_sky6066-1IP | untagged |
| s_sky6066-2IP | untagged |
| s_sky6822-1IP | *Fun30myc sae2Δ* |
| s_sky6822-2IP | *Fun30myc sae2Δ* |
| s_sky7231-1IP | *Fun30myc spo11yf sae2Δ* |
| s_sky7231-2IP | *Fun30myc spo11yf sae2Δ* |
